# Alanine and glutamate catabolism ensure proper sporulation by preventing premature germination and providing energy respectively

**DOI:** 10.1101/2024.03.04.583258

**Authors:** Fengzhi Lyu, Tianyu Zhang, Dong Yang, Lei Rao, Xiaojun Liao

## Abstract

Sporulation as a typical bacterial differentiation process has been studied for decades. However, two crucial aspects of sporulation, (i) the energy sources supporting the process, and (ii) the maintenance of spore dormancy throughout sporulation, are scarcely explored. Here, we reported the crucial role of RocG-mediated glutamate catabolism in regulating mother cell lysis, a critical step for successful sporulation, likely by providing energy metabolite ATP. Notably, *rocG* overexpression resulted in an excessive ATP accumulation in sporulating cells, leading to adverse effects on future spore properties, e.g. increased germination efficiency, reduced DPA content, and lowered heat resistance. Additionally, we revealed that Ald-mediated alanine metabolism decreased the typical germinant L-alanine concentration in sporulating environment, thereby preventing premature germination and maintaining spore dormancy. Our data inferred that sporulation was a highly orchestrated biological process requiring a delicate balance in diverse metabolic pathways, hence ensuring both the completion of sporulation and production of high-quality spores.

## Introduction

Spores are generated by spore-forming bacteria such as the orders *Bacillales* and *Clostridiales* in response to unfavorable environmental conditions, such as nutrient limitation (1, 2). Spores are metabolically dormant and considered the most resilient living organisms due to their extreme resistance to harsh environments, and they are capable of surviving for millions of years (3–5). The process of forming spores from bacterial vegetative cells is termed sporulation. Taking the model bacterium *Bacillus subtilis* as an example, the morphological process of sporulation can be divided into several stages: asymmetric division, engulfment, spore maturation, mother cell lysis, and spore release (6, 7). Initially, as vegetative cells commit to sporulation, the earliest visible event is asymmetric division, which produces the septum, dividing the vegetative cell into a larger mother cell and a smaller forespore. Subsequently, the mother cell membrane migrates around the forespore until it is completely enclosed. This phagocytosis-like process is identified as engulfment. Concurrently, the double-membrane structure of the forespore forms, followed by cortex synthesis and spore coat assembly. Then, the forespore chromosomes become saturated with small acid-soluble proteins (SASPs), and the water within the forespore is replaced by dipicolinic acid (DPA) synthesized in the mother cell, resulting in forespore dehydration. These events culminate in the appearance of phase-bright spores. Next, mother cell lysis occurs after spore maturation, allowing the spores to be released into the environment.

Research on the morphological events of the sporulation process, as well as the underlying gene expression and molecular mechanisms, has been continued for decades (6, 8–11). However, relative few studies focus on guaranteeing proper sporulation, especially concerning the quantity and quality of spores. Proper sporulation encompasses two essential aspects. The first is the normal progression of sporulation. Sporulation is recognized as an energy-intensive biological process (12), implying the importance of energy supply in promoting its progress. Studies have suggested that amino acid metabolism, such as glutamate and alanine catabolism, may serve as potential energy sources driving sporulation (13–15). However, the mechanisms regulating these processes during sporulation remain unclear. Secondly, proper sporulation needs the maintenance of spore dormancy throughout the process. Previous studies report that the deletion of *ylbJ*, *pdaB*, or genes encoding SpoVA protein results in premature germination, indicating the loss of dormancy maintenance ability of generated spores during sporulation (16, 17). The reasons for these types of premature germination are varied, including inappropriate activation of germination receptors, incorrect assembly of spore outer structures, and deficiencies in the SpoVA channel (16, 17). Except for these intrinsic factors, quantities of external factors such as nutrient germinants that induce germination may also exist around the generated spores. How spores maintain dormant in such a tempting environment remains a mystery to be explored. Therefore, defects in either energy supply or dormancy maintenance can lead to abnormal sporulation, adversely affecting the quantity or quality of spores produced.

Here, we reported that the RocG-mediated glutamate catabolism played a crucial role in ensuring proper sporulation, particularly by promoting mother cell lysis through providing energy support. Our research further demonstrated that overexpression of *rocG* resulted in excessively high ATP contents in sporulating cells, which adversely affected the properties of the resulting spores, e.g. elevated germination efficiency, reduced DPA content, and lowered heat resistance. Moreover, we revealed that Ald-mediated alanine catabolism decreased the concentration of typical germinant L-alanine in the sporulating environment to a certain level. This regulation effectively prevented premature germination and contributed to maintain spore dormancy throughout the sporulation process.

## Results

### Proteins involved in alanine, aspartate and glutamate metabolism show enrichment according to proteomics analysis during sporulation of *B. subtilis*

In order to explore the essential metabolic pathways in *B. subtilis* sporulation, Tandem Mass Tag-based (TMT) quantitative proteomics analysis was conducted comparing dormant spores (DS) and sporulating vegetative cells (VC) at t_0_ (Figure 1A). Hierarchical clustering analysis (HCA) was employed to illustrate the overall differences in proteins between the DS and VC groups (Figure 1B). The HCA heatmap displayed greater differences between groups than within groups, indicating significant differences in proteins between DS and VC. Differentially-expressed proteins were identified based on the criteria of p < 0.05 and fold change > 1.2 (the expression level increased by more than 1.2-fold or decreased by less than 0.83-fold). 1,259 proteins with decreased expression as well as 1,248 proteins with increased expression were screened out in the DS group (Figure 1C). KEGG pathway enrichment analysis of these differentially expressed proteins revealed the significant alterations in several metabolic pathways between the DS and VC groups, including alanine, aspartate and glutamate metabolism, ribosome, flagellar assembly, glyoxylate and dicarboxylate metabolism, and methane metabolism (Figure 1D). Of these pathways, the most significant changes were observed in alanine, aspartate and glutamate metabolism, implying their close association with the sporulation of *B. subtilis*. Previous studies have indicated that *ald*, encoding alanine dehydrogenase Ald, and *rocG*, encoding glutamate dehydrogenase RocG, are crucial regulators of alanine and glutamate metabolism, respectively (14, 18). In addition, Δ*ald* and Δ*rocG* mutants have shown notable defects in sporulation (14, 15). However, the deletion of *ansB* gene, encoding L-aspartase important for aspartate metabolism, has no significant effect on sporulation (19). Consequently, further exploration was conducted in the following work to elucidate the effects of alanine and glutamate metabolism on sporulation.

**Figure 1.**
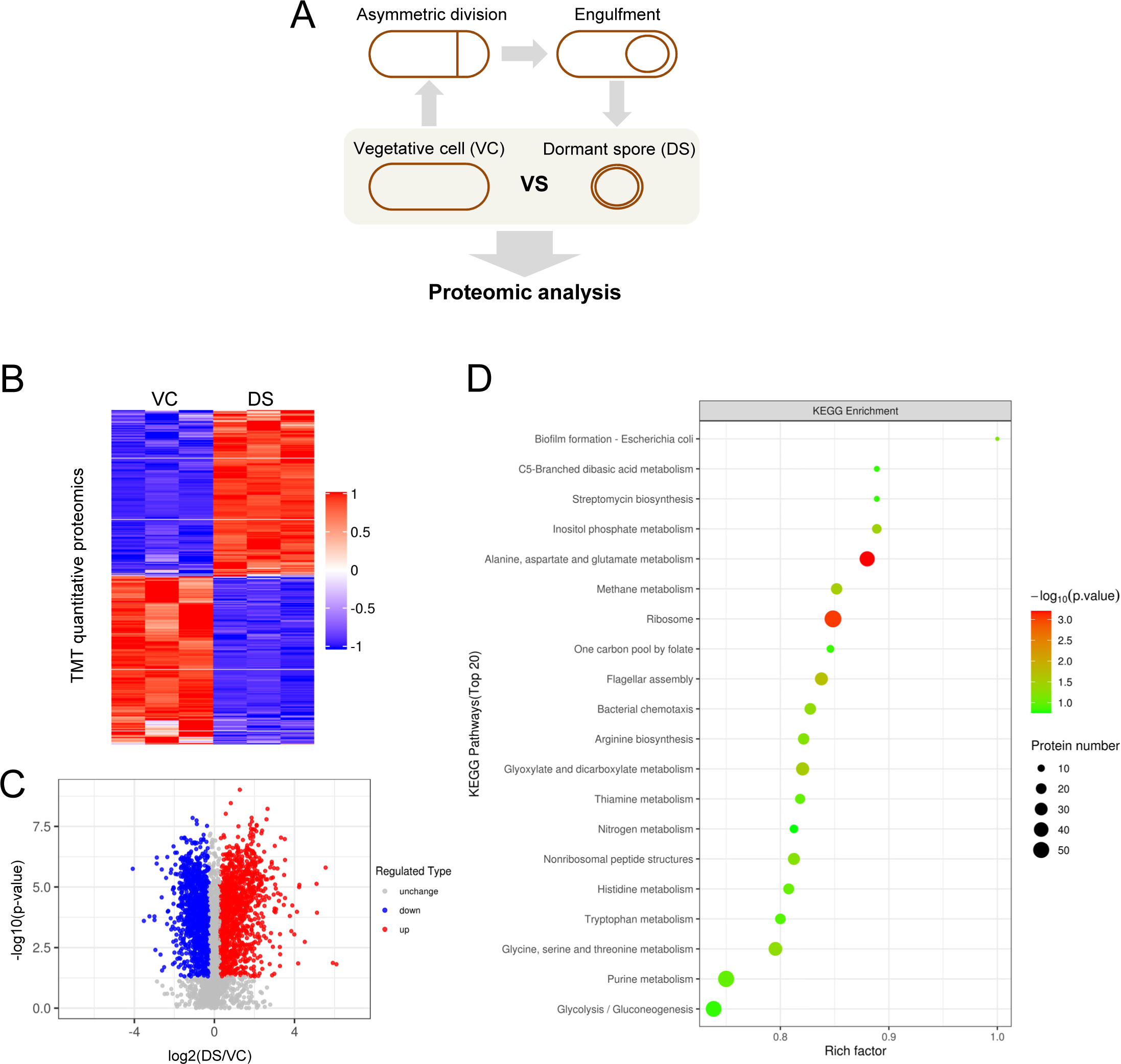
TMT quantitative proteomic analysis. (A) Proteins were compared between dormant spores (DS) and vegetative cells (VC) of at the onset of sporulation (t_0_); (B) Heatmap of differential expressed proteins in DS samples were grouped using Hierarchical Cluster Analysis. Each line represented a protein, with fold change (FC) > 1.2 and p < 0.05 (T-test) as the screening criteria. The proteins with significantly decreased expression were marked in blue, the proteins with significantly increased expression were in red, the proteins without quantitative information were in gray; (C) The volcano map of proteins in DS group was drawn based on FC and the p-value of T test. The proteins with significantly decreased (FC < 0.83, p < 0.05) and increased (FC > 1.2, p < 0.05) expression were marked in blue and red, respectively, and the non-differentiated proteins were in gray; (D) The enrichment map (Top 20) of KEGG pathway enrichment analysis of differentially expressed proteins in the DS group by Fisher’s exact test. The color of the bubble represents the significance of the enriched KEGG pathway, and the color gradient represents the size of the p-value (-log10), and the closer to red, the smaller the p-value. The size of the bubble represents the amount of differential protein.

### Alanine and glutamate metabolism collaboratively regulate sporulation with an additive effect

Given that alanine and glutamate metabolism have been reported separately to be involved in sporulation (14, 15, 20), we wonder if there are any potential joint impacts of these two pathways on sporulation. We constructed the Δ*ald* Δ*rocG* mutant and observed a significantly lower number of phase-bright spores compared to the Δ*ald* or Δ*rocG* mutants (Figure 2A). This indicated that the sporulation defect in the Δ*ald* Δ*rocG* mutant was more pronounced than in either the Δ*ald* or Δ*rocG* mutants. This was further demonstrated by examining the heat-resistant spores produced in sporulation, as the percentage of the Δ*ald* and Δ*rocG* spores was 10.9% and 29.8% respectively, while the Δ*ald* Δ*rocG* mutant was only 0.3% (Figure 2B). The severe sporulation defect of the double mutant suggested that Ald and RocG jointly regulated sporulation in an additive manner. Additionally, we observed an accumulation of phase-dark forespores in the Δ*ald* Δ*rocG* mutant (Figure 2A), which could be attributed to two factors: (i) premature germination due to abnormal spore structure assembly or an inappropriate in-situ sporulating environment (16), or (ii) failure of spore maturation due to limited energy supporting sporulation progression. We first deleted *gerAA* to investigate if premature germination occurred in the Δ*ald* Δ*rocG* mutant. As shown in Figure 3A, the sporulation defect of the double mutant was partially rescued, with 29.7% of phase-bright spores formed in the Δ*ald* Δ*rocG* Δ*gerAA* (Δ3) mutant. This result indicated that premature germination indeed existed in the Δ*ald* Δ*rocG* mutant. However, the partial rescue effect suggested that premature germination was not the exclusive reason for sporulation defect in the Δ*ald* Δ*rocG* mutant. Therefore, the limitation of energy support remained a substantial explanation for the sporulation defect phenotype observed in the double mutant strain. Nonetheless, we further explored these two possibilities in the following work.

**Figure 2.**
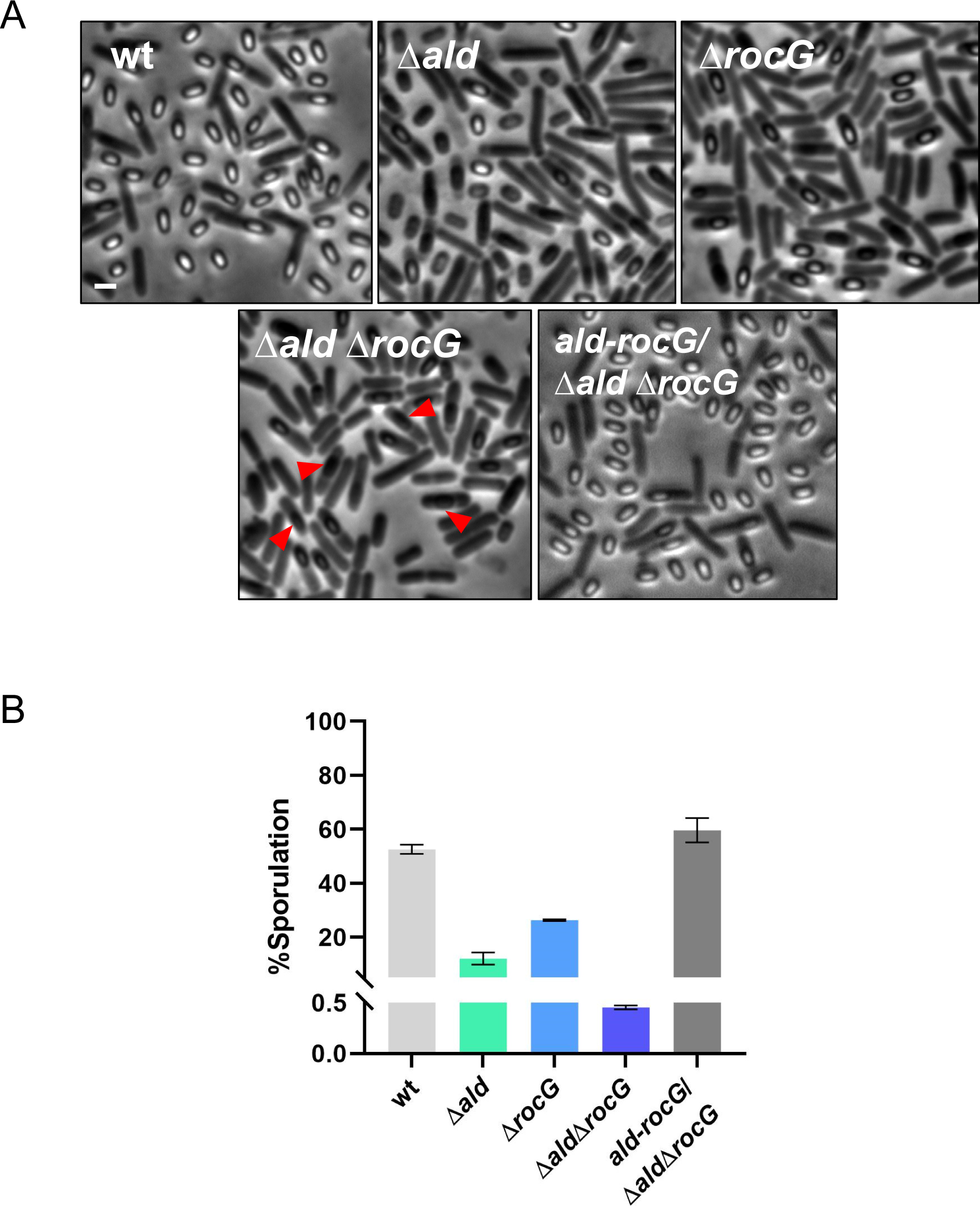
Alanine and glutamate metabolism collaboratively regulate sporulation with an additive effect. (A) Representative phase-contrast images of sporulating cells at the late sporulation stage t_19_. *B. subtilis* PY79 (wt), YZ11 (Δ*ald*), YZ19 (Δ*rocG*), YZ12 (Δ*ald* Δ*rocG*), and YZ13 (Δ*ald* Δ*rocG*, *amyE::ald-rocG*) strains were induced to sporulate in DSM at 37°C for 22 hrs and followed by microscopy. Red arrowheads point to premature germinated spores. Scale bar, 2 μm; (B) The percentage of sporulation of the strains described in (A). Data are presented as the percentage of total number of colonies forming units (CFU) before and after heat treatment (80°C, 20 min). Shown is a representative experiment out of three independent biological repeats.

**Figure 3.**
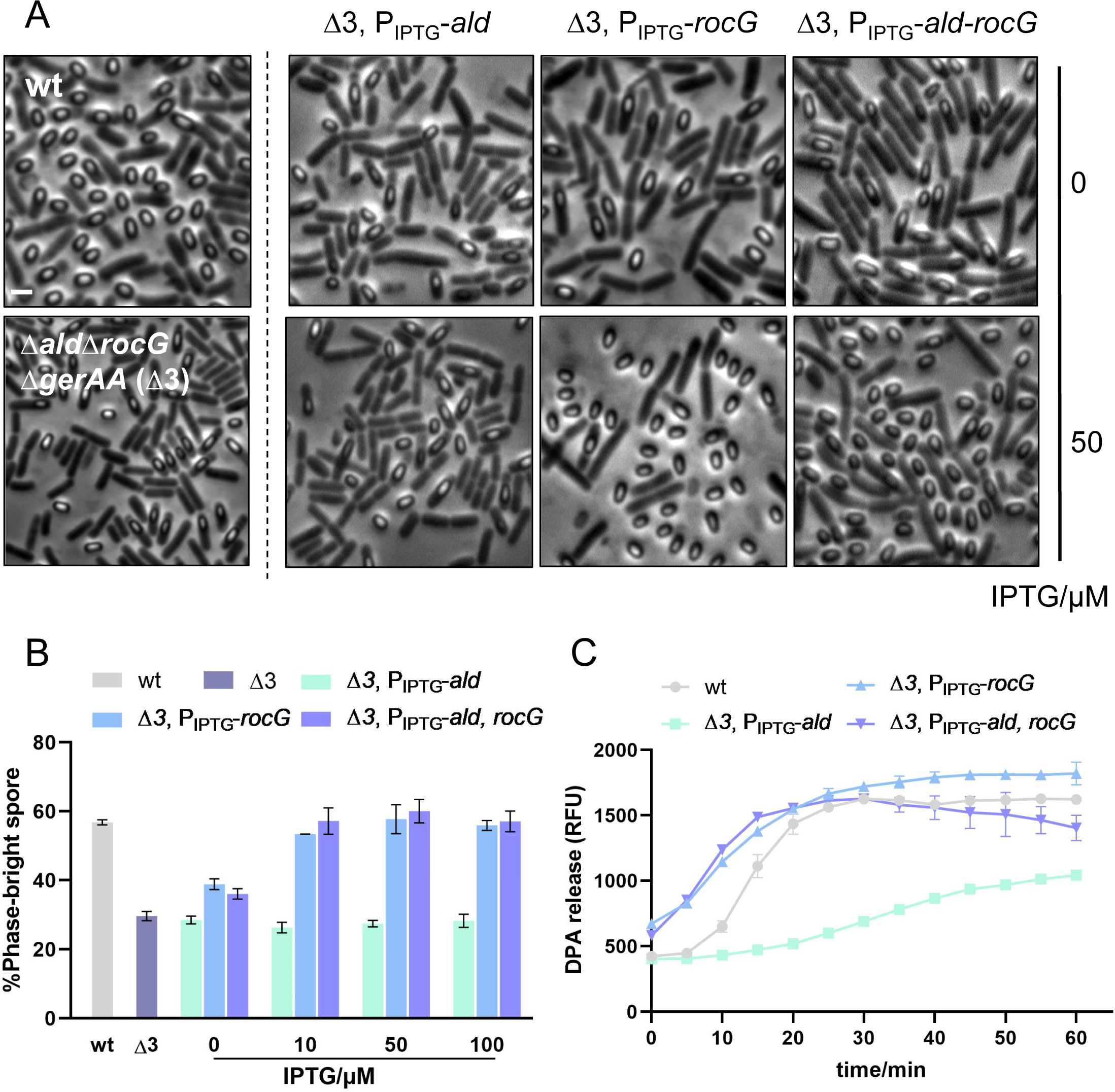
RocG-mediated glutamate metabolism is essential for ensuring the sporulation efficiency. (A) Representative phase-contrast images of sporulating cells at the late sporulation stage t_19_. *B. subtilis* PY79 (wt), YZ22 (Δ*ald* Δ*rocG* Δ*gerAA*), YZ24 (Δ*ald* Δ*rocG* Δ*gerAA, amyE::P_IPTG_-rocG*), YZ25 (Δ*ald* Δ*rocG* Δ*gerAA, amyE::P_IPTG_-ald*), and YZ26 (Δ*ald* Δ*rocG* Δ*gerAA, amyE::P_IPTG_-rocG-ald*) strains were induced to sporulate in DSM at 37°C for 22 hrs and followed by microscopy. 50 μM IPTG was added to the YZ24, YZ25 and YZ26 cultures at the sporulation t_0_ to induce corresponding gene expression. Scale bar, 2 μm; (B) Quantification of the experiment described in (A). Data are presented as percentages of the number of the phase-bright spores and all sporulating cells in the same image (n ≥ 800 for each strain); (C) AGFK-induced germination of spores collected in (A). Spores of wt, as well as YZ24, YZ25, and YZ26 strains with 50 μM IPTG induction, were purified and incubated with AGFK (10 mM) to trigger germination. DPA release was measured by detecting the relative fluorescence units (RFU) of Tb^3+^-DPA. Shown is a representative experiment out of three independent biological repeats.

### RocG-mediated glutamate metabolism, rather than Ald-mediated alanine metabolism, is essential for ensuring both the sporulation efficiency and the spore quality, likely through energy supply

As the Ald and RocG mediated alanine and glutamate metabolisms were proposed as potential energy sources for sporulation (14, 15), we hypothesized that these two metabolic pathways contribute to energy support independently. If this was true, excessive complementation of either metabolic pathway should be capable of rescuing the sporulation defect observed in the Δ*ald* Δ*rocG* mutant. Here, we used the Δ*ald* Δ*rocG* Δ*gerAA* (Δ3) strain to exclude the premature germination effect and independently explored the energy supply mechanism (Figure 3A). Based on this, *ald* and *rocG* were artificially expressed separately and jointly in Δ3 under an IPTG-inducible promoter. Results indicated that increasing the expression level of *ald* by raising the concentration of IPTG up to 5 mM had no significant effect on the quantity of phase-bright spores in the Δ3 mutant, with the sporulation percentage ranging between 20% and 30% (Figure S1, Figure S2A). Accordingly, these spores exhibited significant germination deficiency under AGFK induction (Figure S2B, Figure 3C). However, elevating the *rocG* expression level with the addition of at least 10 mM IPTG restored the sporulation of the Δ3 mutant to 53.4%, similar to that of the wild-type (56.8%) (Figure 3B). Furthermore, when more than 20 mM IPTG was added, the germination deficiency of the Δ3 mutant spores was also recovered to the level of the wild-type (Figure S2B, Figure 3C). Notably, the Δ3 mutant with IPTG-induced co-expression of *ald* and *rocG* exhibited the same sporulation and germination phenotypes as those with IPTG-induced sole expression of *rocG* (Figure S2B, Figure 3C). Hence, in the Δ3 mutant, sole complementation of RocG succeeded in rescuing the sporulation defect, unlike in the case of Ald. This indicated that RocG-mediated glutamate metabolism appeared to regulate sporulation by providing energy sources, whereas Ald-mediated alanine metabolism may not play the same role.

To further investigate whether these two catabolic pathways are involved in controlling the spore quality, mutants of transcription factors of Ald and RocG, Δ*adeR* and Δ*ahrC* Δ*rocR* (15, 21), were respectively constructed to examine the germination phenotype of the spores. *gerAA* was also knocked out in these mutants to ensure the comparability of experimental results. The deletion of transcription factors was demonstrated to reduce, rather than completely eliminate, the expression of regulated genes (Figure 4B), and this rescued the sporulation deficiency (Figure 4A). Interestingly, no significant germination defect was observed in spores of the Δ*adeR* Δ*gerAA* mutant with decreased expression of *ald* (Figure 4C). However, spores of the Δ*ahrC* Δ*rocR* Δ*gerAA* mutant with low expression of *rocG* showed a remarkable germination deficiency (Figure 4C). Moreover, spores of the Δ*adeR* Δ*ahrC* Δ*rocR* Δ*gerAA* mutant exhibited a similar germination deficiency phenotype to that of Δ*ahrC* Δ*rocR* Δ*gerAA* mutant spores (Figure 4C), indicating that the expression of *rocG*, rather than *ald*, was essential for ensuring spore quality with normal germination capability. Taken together, these results strongly implied that RocG-mediated glutamate metabolism regulated both sporulation efficiency and spore quality, probably through energy supply. As for Ald-mediated alanine metabolism, its effect on sporulation was unlikely to be executed by providing energy sources. Instead, it is more reasonably associated with premature germination, as quantities of phase-dark spores were observed in the sporulating cells of the Δ*ald* mutant (Figure 2A).

**Figure 4.**
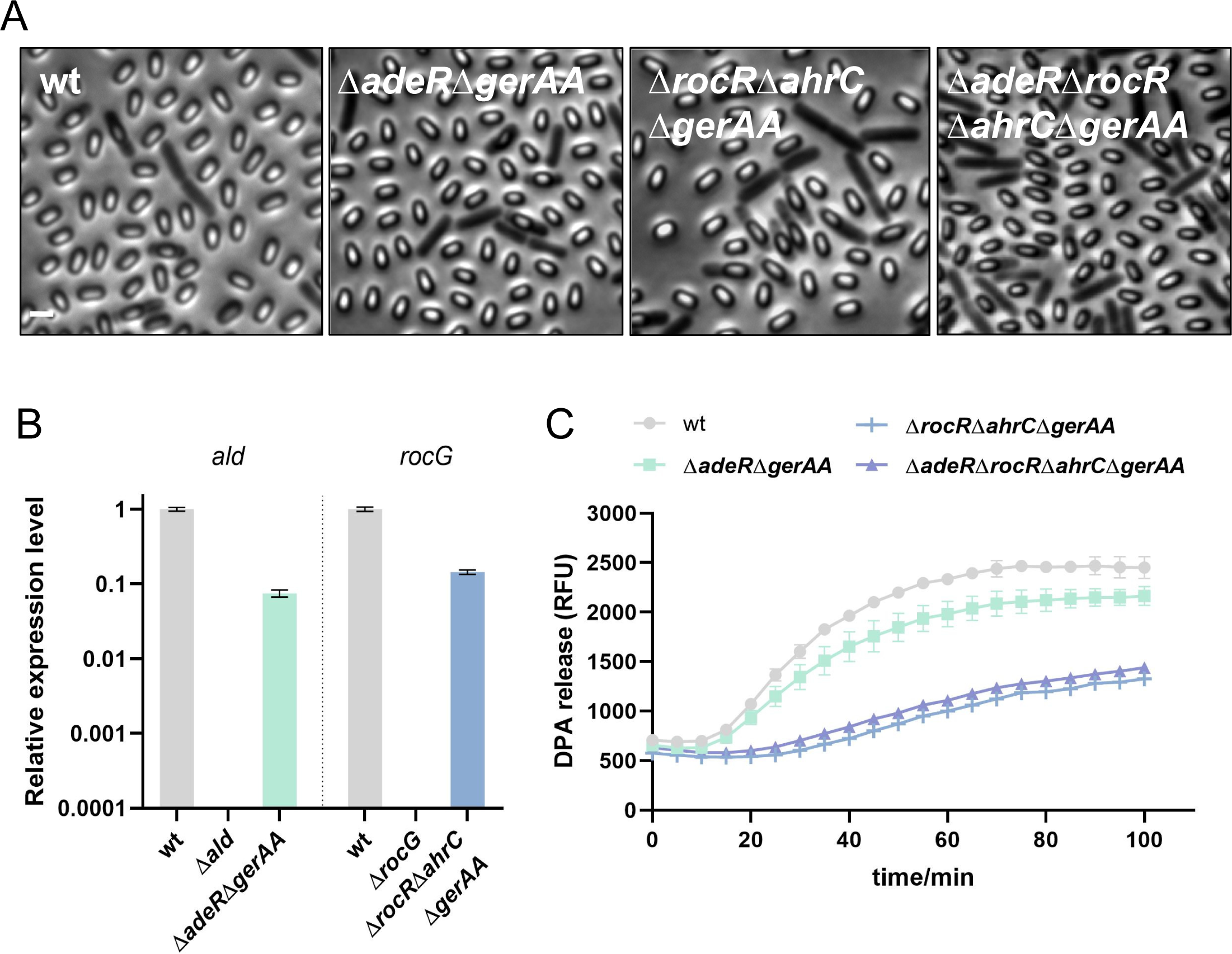
The expression of *rocG* during sporulation is crucial to ensure the quality of correspondingly generated spores. (A) Representative phase-contrast images of sporulating cells at the late sporulation stage t_19_. *B. subtilis* PY79 (wt), YZ81 (Δ*adeR* Δ*gerAA*), YZ23 (Δ*ahrC* Δ*rocR* Δ*gerAA*), and YZ90 (Δ*adeR* Δ*ahrC* Δ*rocR* Δ*gerAA*) strains were induced to sporulate in DSM at 37°C for 22 hrs and followed by microscopy. Scale bar, 2 μm; (B) Expression of the *ald* gene in wt, YZ11 (Δ*ald*) and YZ81 strains, and *rocG* gene in wt, YZ19 (Δ*rocG*), and YZ23 strains. Sporulating cells were collected at t_0_ and detected as described in Methods; (C) AGFK-induced germination of spores collected in (A). Spores of wt, YZ81, YZ23, and YZ90 strains were purified and incubated with AGFK (10 mM) to trigger germination. DPA release was measured by detecting the RFU of Tb^3+^-DPA. Shown is a representative experiment out of three independent biological repeats.

### Ald inhibits premature germination during sporulation by reducing L-alanine content in the external environment of generating spores

As mentioned above, the presence of phase-dark spores in the Δ*ald* mutant prompted us to investigate the impact of Ald-mediated alanine metabolism on premature germination. As illustrated in Figure 5A-5B, the Δ*ald* mutant produced significant quantities of phase-dark spores, and the percentage of phase-bright spores was only 11.5%. However, upon the deletion of *gerAA* in the Δ*ald* mutant, the percentage of phase-bright spores significantly raised to 61.5%, approaching levels observed in the wild-type (76.5%). In addition, no notable deficiency in germination was observed in Δ*ald* Δ*gerAA* spores compared to the wild-type (Figure 5C). Thus, it can be concluded that the absence of Ald led to sporulation defect by inducing premature germination during sporulation. To identify the timing of premature germination occurrence, the sporulation process of the Δ*ald* mutant was examined by time-lapse microscopy. Interestingly, two models were observed: (i) forespores prematurely germinated during mother cell lysis, and then were released; (ii) dormant spores were released and subsequently induced to premature germination (Figure 5D).

**Figure 5.**
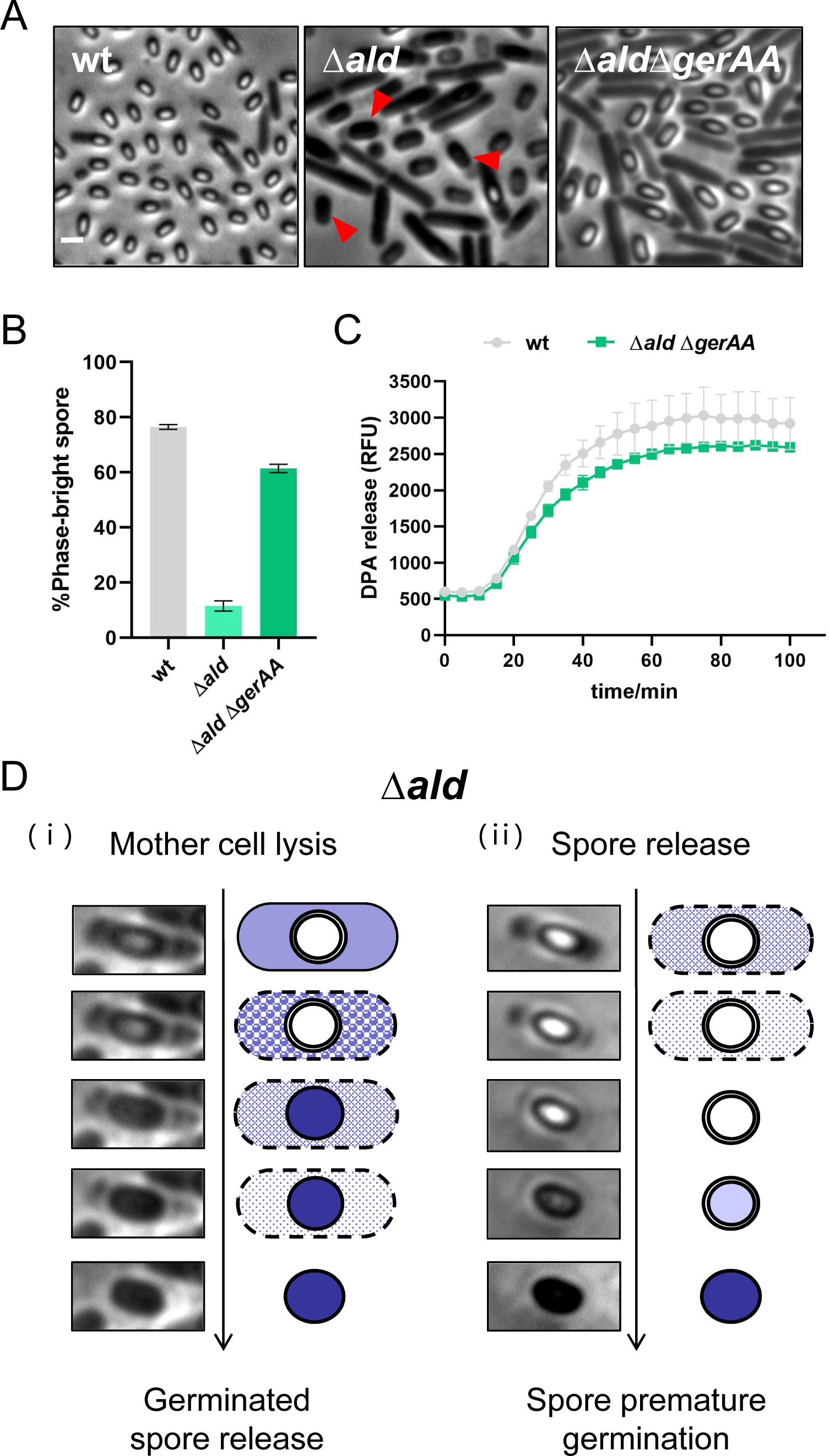
The absence of Ald induces premature germination during sporulation. (A) Representative phase-contrast images of sporulating cells at the late sporulation stage t_19_. *B. subtilis* PY79 (wt), YZ11 (Δ*ald*), and YZ21 (Δ*ald* Δ*gerAA*) strains were induced to sporulate in DSM at 37°C for 22 hrs and followed by microscopy. Red arrowheads point to premature germinated spores. Scale bar, 2 μm; (B) Quantification of the experiment described in (A). Data are presented as percentages of the number of the phase-bright spores and all sporulating cells in the same image (n ≥ 800 for each strain); (C) AGFK-induced germination of spores collected in (A). Spores of wt, YZ11, and YZ21 strains were purified and incubated with AGFK (10 mM) to trigger germination. DPA release was measured by detecting the RFU of Tb^3+^-DPA. Shown is a representative experiment out of three independent biological repeats; (D) Models of the premature germination in the Δ*ald* mutant. YZ11 strain was induced to sporulate in DSM at 37°C. After 14 hrs of incubation, sporulating cells were collected on an exhausted DSM gel-pad as described in Methods, and followed by time-lapse microscopy at a 10 min interval.

The connection between Ald and premature germination raised up an intriguing speculation that the interruption of alanine metabolism may lead to an over-accumulation of L-alanine, which triggers premature germination. To investigate this, *ald* was artificially expressed in the Δ*ald* mutant under an IPTG-inducible promoter, and the sporulation phenotype as well as the environmental L-alanine content at the late sporulation phase (t_19_) of these mutants were examined. The results showed a gradual increase in the percentage of phase-bright spores with elevating the expression levels of *ald* (Figure 6A-6C). Notably, the expression of *ald* decreased as the concentration of added IPTG increased to 1 mM, which was possibly due to the toxicity of IPTG to cells (Figure 6C). In contrast, an opposite trend was observed in the environmental L-alanine concentration of the Δ*ald* mutant, which reached 3,397.4 μM without IPTG induction, while decreased to wild-type levels of 145.9 μM when > 200 μM IPTG was added to elevate the *ald* expression (Figure 6B). Consequently, Ald-mediated alanine metabolism was responsible for reducing the L-alanine content in the external environment of spores to prevent premature germination, and thus ensuring proper sporulation.

**Figure 6.**
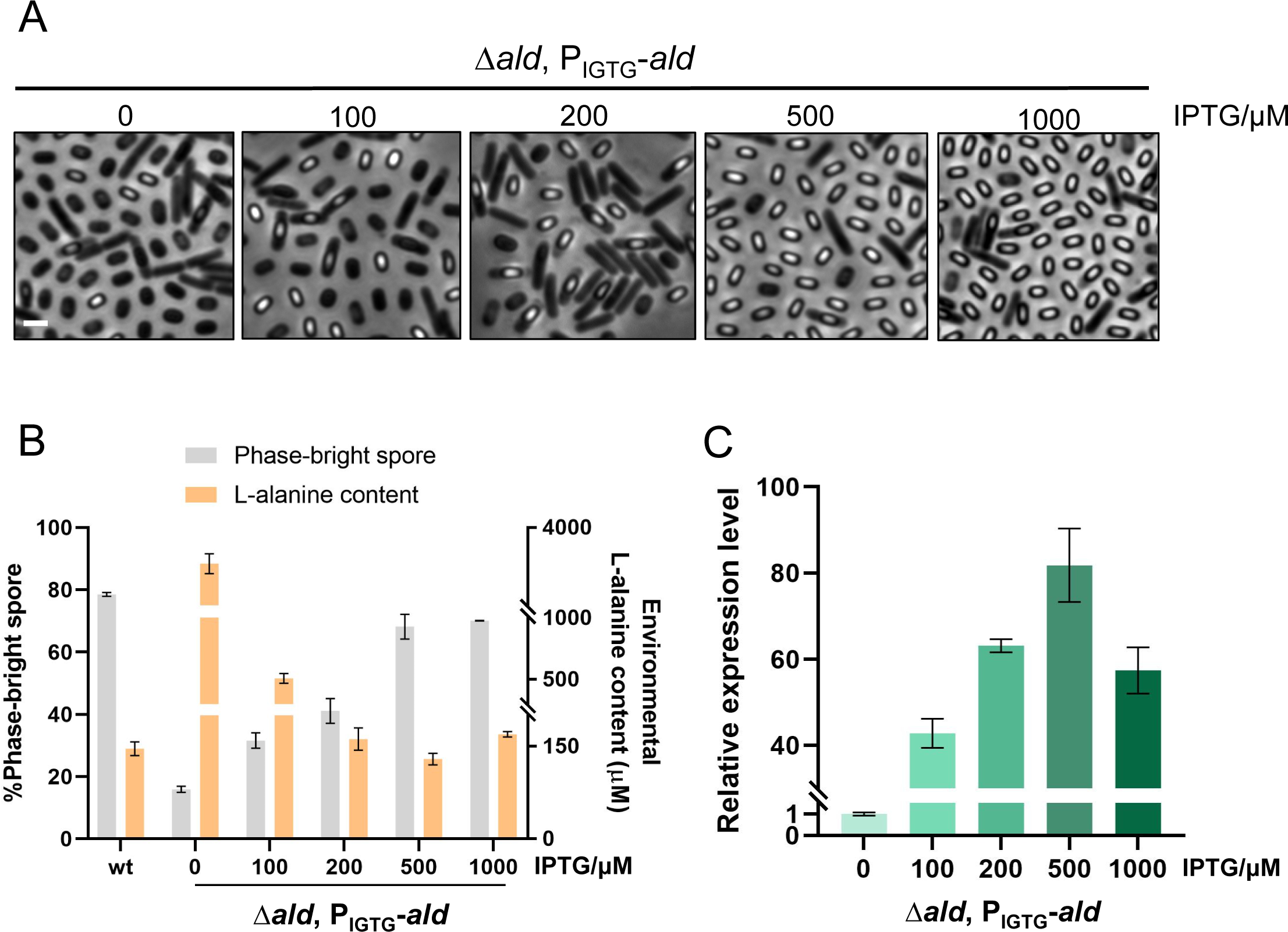
Ald-mediated alanine metabolism regulates L-alanine content in the external environment of generated spores. (A) Representative phase-contrast images of sporulating cells at the late sporulation stage t_19_. YZ31 (Δ*ald, amyE::P_IPTG_-ald*) strains were induced to sporulate in DSM at 37°C for 22 hrs and followed by microscopy. 0-1000 μM IPTG was added at the sporulation t_0_ to induce *ald* expression. Scale bar, 2 μm; (B) The percentage of phase-bright spores as well as the environmental L-alanine content of the wt and YZ31 strains at the late sporulation stage t_19_. The percentage of phase-bright spores are presented as ratio of the number of the phase-bright spores and all sporulating cells in the same image (n ≥ 800 for each strain); The environmental L-alanine content was detected as described in Methods; (C) Expression of the *ald* gene in YZ31 strains with different IPTG induction. Sporulating cells were collected 10 min after IPTG addition and detected as described in Methods. Shown is a representative experiment out of three independent biological repeats.

### RocG plays a crucial role in regulating both σ^K^-dependent spore release and spore properties likely by providing energy sources

As indicated above, RocG-mediated glutamate metabolism regulated both sporulation efficiency and spore quality likely through energy support. To further explore the underlying mechanism, we focused on identifying the specific sporulation stage interrupted by *rocG* deletion. We constructed strains capable of reporting stage-specific sigma factors involved in sporulation, namely σ^F^, σ^E^, σ^G^, and σ^K^, which respectively regulated polar division, engulfment, spore maturation, and spore release (6, 11, 22, 23). P*_spoIIQ_*, P*_spoIID_*, P*_sspB_*, and P*_gerE_* promotors that was respectively recognized by σ^F^, σ^E^, σ^G^, and σ^K^ were fused to *gfp*. In general, the activation of a particular σ factor correlates with σ-dependent GFP fluorescence, thereby visually displaying the impaired sporulation stage (22). Our results demonstrated that the activation patterns of σ^F^ and σ^E^ in the Δ*rocG* mutant were similar to those of the wild-type (Figure S3). The activation of σ^G^ could be achieved in the Δ*rocG* mutant, though there was a delay in the activation timepoint (Figure 7A). Remarkably, σ^K^-dependent GFP fluorescence in the Δ*rocG* mutant initially appeared at t_7_ but abnormally persisted until t_25_ (Figure 7B). Since σ^K^ regulates cell wall lytic enzymes, leading to mother cell lysis (24, 25), its fluorescence is supposed to disappear with spore release. The persistence of σ^K^-dependent GFP fluorescence at the end of sporulation in the Δ*rocG* mutant indicated the failure of mother cell lysis, a process mainly executed by the sporulation-specific cell wall lytic enzymes CwlC and CwlH (24, 26). As expected, the expression level of *cwlC* and *cwlH* were remarkably lower in the Δ*rocG* mutant compared to those in the wild-type (Figure 7C). Taken together, the deletion of *rocG* had no notable effect on the activation of sporulation-specific σ factors but hindered mother cell lysis by impacting the expression of cell wall lytic enzymes, resulting in an impaired spore release process (Figure 7D).

**Figure 7.**
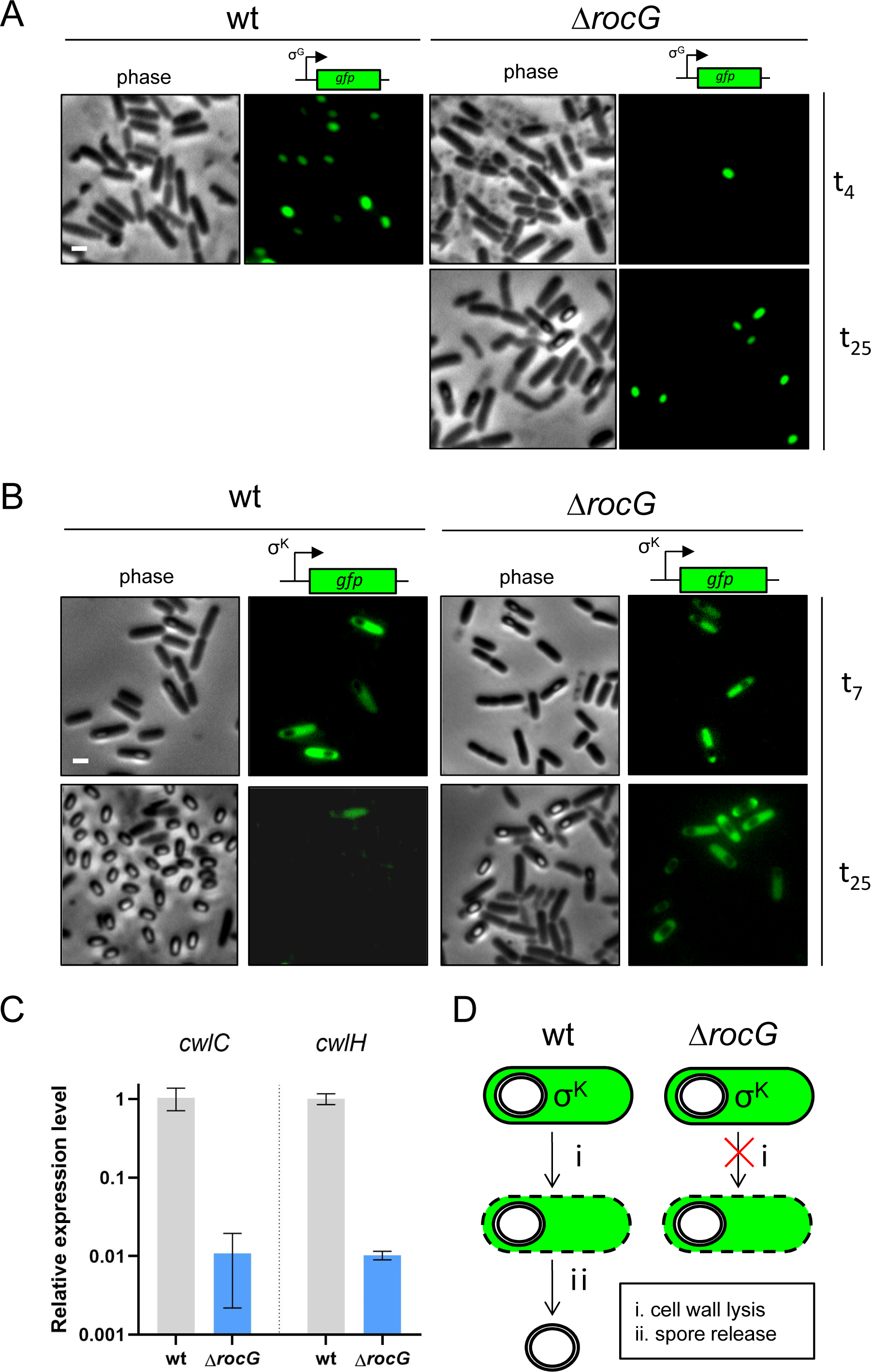
Cytological sporulation assay reveals the impaired sporulation stage of Δ*rocG* mutants. (A-B) Representative phase contrast and the indicated fluorescent images of wt and YZ19 (Δ*rocG*) cells harboring two transcriptional fusions, σ^G^ and σ^K^, at sporulation t_4_, t_7_, and t_25_. Scale bar, 2 μm; (C) Expression of the *cwlC* and *cwlH* gene in wt and YZ19 strains. Sporulating cells were collected at t_7_ and detected as described in Methods. Shown is a representative experiment out of three independent biological repeats. (D) Models of the sporulation defect in the Δ*rocG* mutants.

To further understand the regulating role of RocG in sporulation, *rocG* was artificially expressed in the Δ*rocG* mutant under an IPTG-inducible promoter. The results showed a significant increase in the percentage of released spores with elevating expression levels of *rocG* (Figure 8A-8C). Moreover, the addition of more than 10 μM IPTG remarkably improved the percentage of spore release to the wild-type level (Figure 8A, 8C). As hypothesized previously, RocG-mediated glutamate metabolism could provide energy sources to drive sporulation proceeding, we then examined the ATP levels in sporulating cells to verify this. Since previous research has demonstrated that the ATP content of mother cells during sporulation peaked at t_1_ (15, 27), we examined ATP content at t_1_ in the Δ*rocG* mutant with different *rocG* expression levels. Accordingly, the level of ATP in the Δ*rocG* mutant increased with the elevation of *rocG* expression (Figure 8D). The positive correlation between the ATP level in sporulating cells (Figure 8D) and future spore release (Figure 8C) suggested the crucial role of energy supply in the late sporulation process, particularly in mother cell lysis. Notably, the ATP content of the Δ*rocG* mutant with 50 μM IPTG induction was almost double that of the wild-type (Figure 8D). We then wondered if such a high level of ATP in sporulating cells could affect the properties of the future spores. To test this, the spores generated under different concentrations of IPTG induction were purified and examined for germination phenotypes, as well as DPA content and heat resistance. Interestingly, Δ*rocG* spores with 50 μM IPTG induction showed higher germination efficiency (Figure 8G-8I) but significantly lower DPA content (Figure 8E) as well as decreased heat resistance (Figure 8F) compared to the wild-type spores. Taken together, the expression of *rocG* can indeed provide energy support for sporulation, especially for mother cell lysis regulated by σ^K^, thus contributing to the proper sporulation process. However, overexpression of *rocG* can accumulate excessive ATP in sporulating cells, which might adversely affect the spore properties.

**Figure 8.**
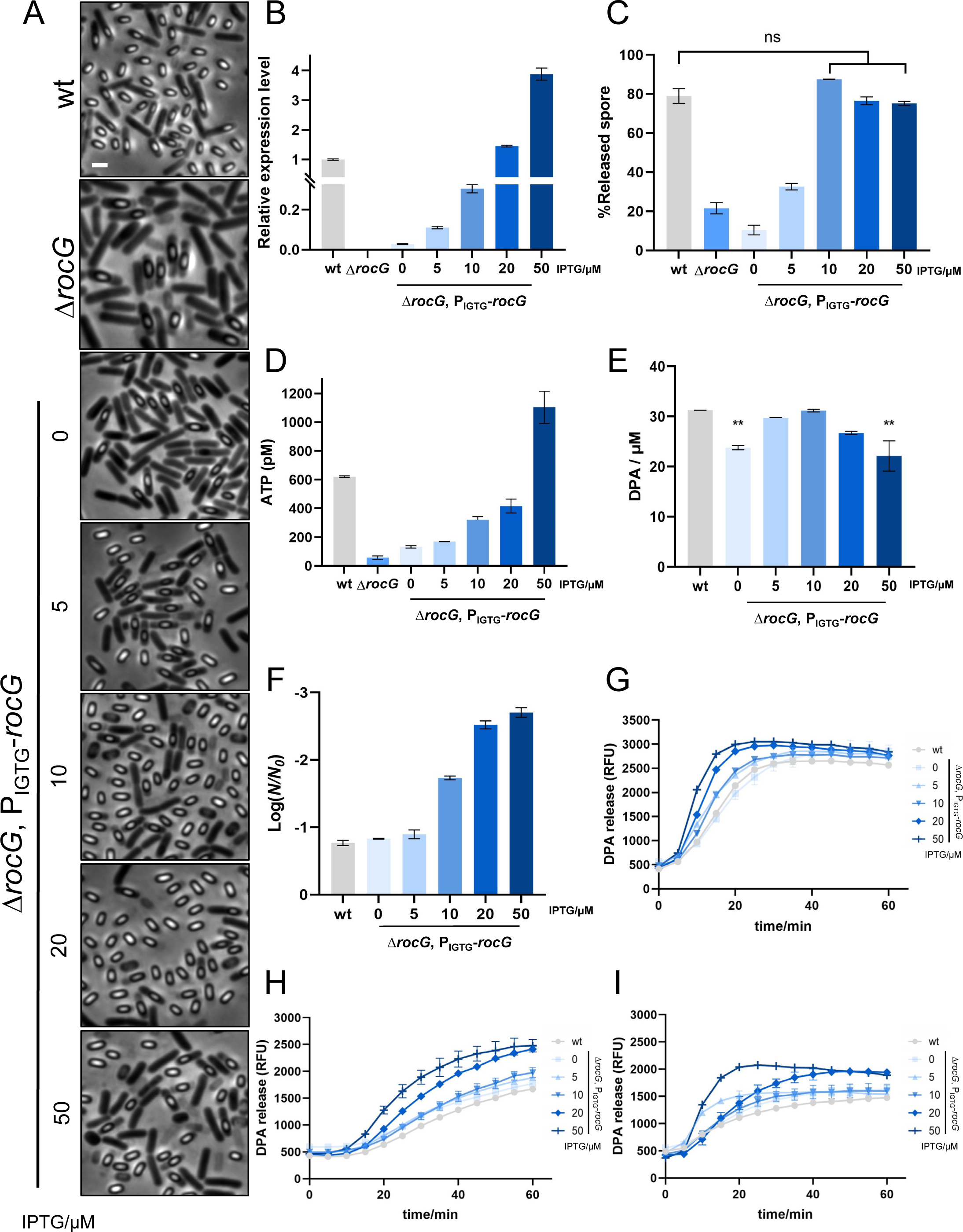
RocG regulates both mother cell lysis and spore properties. (A) Representative phase-contrast images of sporulating cells at the late sporulation stage t_19_. *B. subtilis* PY79 (wt), YZ19 (Δ*rocG*) and YZ32 (Δ*rocG, amyE::P_IPTG_-rocG*) strains were induced to sporulate in DSM at 37°C for 22 hrs and followed by microscopy. 0-50 μM IPTG was added to YZ32 at the sporulation t_0_ to induce *rocG* expression. Scale bar, 2 μm; (B) Expression of the *rocG* gene in strains indicated in (A). Sporulating cells were collected 30 min after IPTG addition and detected as described in Methods; (C) Quantification of released spores produced by strains indicated in (A). Data are presented as percentages of the number of the released spores and all sporulating cells in the same image (n ≥ 800 for each strain); (D) ATP levels in strains indicated in (A). Sporulating culture was collected at t_1_ and analyzed for ATP level as described in Methods; (E) DPA content in spores of wt and YZ32 with different IPTG induction. Spores were purified and boiling for 20 min. DPA content was measured by detecting the RFU of Tb^3+^-DPA; (F) Heat resistance of spores collected in (E). Data are presented as the percentage of total number of CFU before and after heat treatment (90°C, 10 min); (G-I) Germination phenotypes of spores collected in (E). Spores were purified and incubated with (G) L-alanine (10 mM), (H) AGFK (10 mM), and (I) DDA (10 mM) to trigger germination. DPA release was measured by detecting the RFU of Tb^3+^-DPA. Shown is a representative experiment out of three independent biological repeats.

## Discussion

Sporulation as a typical bacterial differentiation process has been extensively studied for decades, and the morphological events along with the signal transduction for this process are relatively well elucidated (6, 8–11). However, as an energy-consuming process, the sources of energy supply and the underlying regulating mechanism lack research. In addition, how the generated spores maintain in dormant state during sporulation remains mysterious. Here, we demonstrated that Ald-mediated alanine metabolism decreased the concentration of the typical germinant L-alanine in the sporulating environment to a certain level, thus avoiding premature germination and maintaining spore dormancy. Moreover, we also provided evidences supporting that RocG-mediated glutamate metabolism ensured proper sporulation, especially mother cell lysis, by regulating ATP levels during sporulation. Additionally, excessively high ATP levels during the sporulation process was supposed to adversely affect the properties of the produced spores, including faster germination efficiency, lower DPA content, along with decreased heat resistance. Our data revealed that sporulation was a highly orchestrated and exquisite biological process requiring the balance of diverse metabolic pathways, e.g. alanine catabolism to eliminate surrounding germinants, and glutamate metabolism providing an appropriate level of energy to ensure both sporulation completion and the high quality of generated spores (Figure 9).

**Figure 9.**
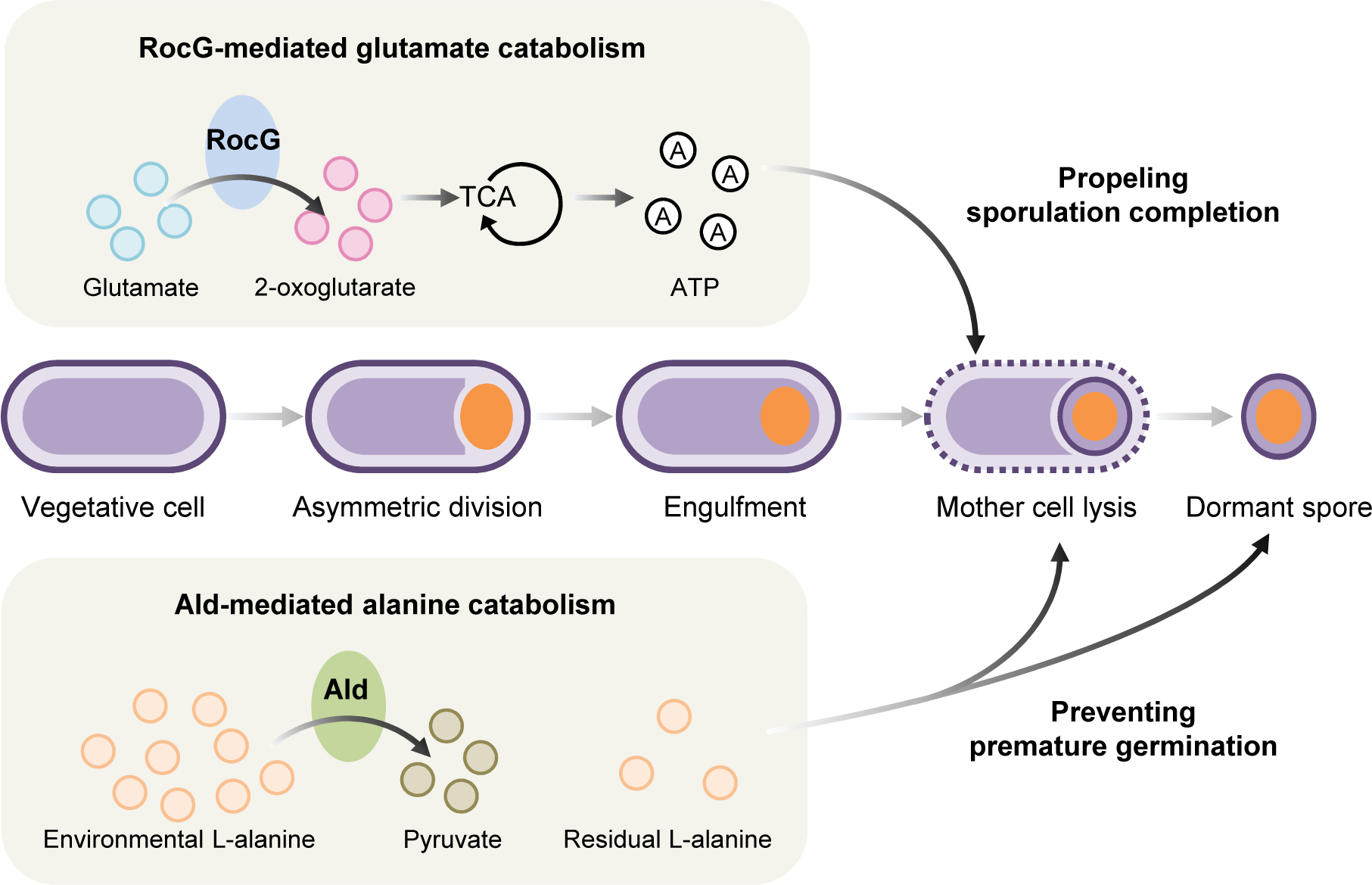
The crucial roles of alanine and glutamate catabolism in ensuring proper sporulation.

Our finding of alanine catabolism eliminating the germinant L-alanine raises another open question that which catabolic pathways or biological reactions are responsible for regulating the balance of other potential germinants during sporulation. Indeed, sporulation involves protein turnover, in which new proteins are continuously synthesized using the amino acids derived from the breakdown of pre-existing cellular protein (28, 29). Hence, substantial free amino acids exist during sporulation, which could serve as potential germinants. Moreover, non-sporulating cells can produce meso-diaminopimelic acid (m-DAP) type muropeptides, also identified as possible germinants (30). However, the mechanisms by which spores eliminate these potential germinants remain unclear. Our research revealed that alanine catabolism is one of the strategies employed to achieve this. While we also observed that L-alanine was not completely eliminated, as 145.9 μM L-alanine was still detected in the sporulating medium of the wild-type (Figure 6B). Why did the presence of residual L-alanine in the environment not trigger germination? One plausible explanation is that alanine racemases present in the spore coat can convert the germinant L-alanine into the germination inhibitor D-alanine, thereby allowing the spores to persist in dormancy (31). However, whether there are alternative mechanisms preventing the residual L-alanine and other germinants from triggering germination is worth exploring.

Another new finding in our study revealed that RocG-mediated glutamate metabolism plays a crucial role as an energy source for sporulation. Indeed, the catabolic product of glutamate, 2-oxoglutarate (2-OG), directly participates in the tricarboxylic acid (TCA) cycle, showing its superior efficiency in providing energy during nutrient-limited sporulation (32, 33). Concurrently, various amino acids such as proline, ornithine, citrulline, and arginine can be converted into glutamate through the arginine degradation pathway (34), indicating its high availability in sporulating cells. These aspects give glutamate an advantage in supporting energy during sporulation compared to other amino acids. Additionally, glutamate stands out among the limited free amino acids found within diverse spores (35), further suggesting its crucial role during sporulation. In our study, we showed that ATP produced by glutamate catabolism is highly corelated with mother cell lysis regulated by the cell wall lytic enzymes CwlC and CwlH, and the interruption of this catabolism remarkably reduced the expression of these two enzymes. This finding implies that the energy derived from glutamate catabolism is crucial for the expression of genes regulating the final stage of sporulation. Moreover, we also observed that the overaccumulation of ATP in sporulating cells through glutamate catabolism adversely affected the properties of future spores. These effects may be attributed to abnormal changes in the structure assembly or molecular reservoir in the generated spores (28, 36). Actually, spores can inherit molecules from sporulating cells, such as alanine dehydrogenase and ATP, to modulate their revival capability (36). Moreover, glutamate, as a universal amino group donor in all living organisms, can serve as a carbon or nitrogen source for synthesizing other amino acids and DPA during sporulation (34, 35, 37). Additionally, the glutamate catabolic intermediate 2-OG can contribute to the synthesis of amino acids, nucleotides, and NADH (32), potentially affecting the structure construction or molecular modulation of spores. All these evidences support that glutamate is an ideal substrate for energy supply and the synthesis of new substances during sporulation, ensuring both the proper sporulation process and the quality of the spores.

## Methods

### Strains and plasmids

*B. subtilis* strains used in this study are listed Table S1. Plasmids construction is listed in Table S2, and primers are described in Table S3. For gene replacement strategy, primer pairs were used to amplify the flanking genomic regions of the corresponding gene. PCR products and the respective antibiotic resistance gene were used for Gibson assembly (NEB, USA) (38). The product was used to transform *B. subtilis* PY79 to obtain the mutant allele.

### General methods

All general methods for *B. subtilis* were carried out as described previously with some modifications (15). Cultures of wild-type and mutant strains were cultivated in LB medium (Difco) at 37 °C. Sporulation was carried out at 37°C by suspending overnight cells (OD_600_ = 0.05) in Schaeffer’s liquid medium (Difco Sporulation Medium, DSM) (39). Sporulation t_0_ was identified as the third hour after spores suspending in DSM. The percentage of sporulation was evaluated by calculating the ratio of total number of colonies forming units (CFU) before and after heat treatment (80°C, 20 min) (40). The percentage of phase-bright or released spores were counted based on the according phase-contrast images. To ensure confidence of the data, > 800 cells were counted for each experiment. Spore germination with different germinants was examined as described previously with some modifications (15, 41). Briefly, purified spores were heat activated at 75°C for 30 min, and then induced by L-Alanine (10 mM) or AGFK (2.5 mM L-Asparagine, 5 mg/mL D-glucose, 5 mg/mL D-fructose, and 50 mM KCl) at 37°C, or DDA (1 mM in 10 mM Tris-HCl, pH 7.4) at 42°C. The germination was tested by determining the DPA release as descried in the following text.

### Spore purification

Matured spores were purified as described previously (15). Briefly, 22 hrs DSM culture was centrifuged and washed 3 times by DDW and then kept in 4°C with constant agitation. The suspension was washed once a day and resuspended in DDW. After 7 days, the suspension was centrifuged to collect the pellet. 20% histodenz solution was used to resuspend the pellet at a ratio of 400 μL per 10 mL of DSM for 30 min on ice. Aliquots (200 μL) of resuspension mixture were then added on top of 900 μL 50% histodenz solution, and gradient fractionation was carried out by centrifugation at 15,000 rpm at 4°C for 30 min. The pellet was collected and washed at least 5 times by DDW. Phase contrast microscopy was then used to evaluate the purity of pellet spores. Spores with >99% purity can be used for following experiments, otherwise the purification steps should be carried out more than once.

### Tandem Mass Tag-based (TMT) quantitative proteomics analysis

TMT quantitative proteomics analysis was carried out between pure dormant spores (DS) and vegetative cells (VC) at sporulation t_0_ by Shanghai Applied Protein Technology Co., Ltd (Shanghai, China). DS and VC samples were collected by centrifugation and then freeze-dried and bead-grinded using FastPrep-24 (M. P. Biomedicals, LLC, USA). Samples were then extracted for proteins and labeled using TMT reagent. Proteomics analysis was then carried out by LC-MS/MS system with on a Q Exactive mass spectrometer (Thermo Scientific) that was coupled to Easy nLC (Proxeon Biosystems, now Thermo Fisher Scientific). The raw data for each sample were searched using the MASCOT engine (Matrix Science, London, UK; version 2.2) embedded into Proteome Discoverer 1.4 software for identification and quantitation analysis. Hierarchical clustering analysis was performed using Cluster 3.0 (http://bonsai.hgc.jp/~mdehoon/software/cluster/software.htm) and Java Treeview software (http://jtreeview.sourceforge.net). Enrichment analysis was performed based on KEGG database (http://geneontology.org/). A statistical analysis was performed using a t-test to determine the significance (p-value) of differentially-expressed proteins. The expression level of proteins with p < 0.05 and fold change > 1.2 (the expression level increased by more than 1.2-fold or decreased by less than 0.83-fold) were considered as significant difference.

### DPA measurements

DPA release was detected as described previously with some modifications (42). Briefly, spore germination was induced by L-Ala, AGFK or DDA at 37°C or 42°C in a 96-well plate. Spores at OD_600_ of 20, 10 mM germinants, 25 mM K-Hepes buffer (pH 7.4) as well as 50 mM TbCl_3_ were mixed in 200 μL and Tb^3+^-DPA fluorescence intensity was monitored at Ex/Em = 270/545 nm by a TECAN Spark 10M microplate reader (TECAN, Switzerland). Total DPA content of spores were evaluated by boiling the spores (OD_600_ of 1) for 20 min and mixing the spores and 50 mM TbCl_3_ to 200 μL in a 96-well plate. The DPA standard solution was serially diluted and detected together to obtain a standard curve. The detection parameters for DPA release were the same as above, and the total DPA content of spores was calculated based on the standard curve.

### Phase-contrast and fluorescence microscopy

Phase-contrast and fluorescence microscopy were performed using a Nikon DS-Qi2 microscope equipped with a Nikon Ph3 DL 100x/1.25 Oil phase contrast objective. Both bacterial cells (500 μL) and spores (50 μL) were centrifuged, and the pellets were resuspended with 5 ∼ 10 μL PBSx1 and then imaged. For time-lapse imaging of sporulation, Imaging System Cell Chamber (AttofluorTM Cell Chamber) was used. Sporulating cells were collected by centrifugation. The supernatant DSM was collected to make a gel-pad with 1% agarose. The collected sporulating cells were incubated on the DSM gel-pad in a chamber at 37 °C. Image analysis and processing were performed by ImageJ2.

### Real-Time Quantitative PCR (RT-qPCR)

Real-time quantitative PCR (RT-qPCR) was carried out followed the protocol described previously (15). RNA samples of sporulating cells (500 μL) were collected from DSM by centrifugation, and then extracted by FastPure Cell/Tissue Total RNA Isolation Kit V2 (Vazyme Biotech Co..Ltd). HiScript III All in-one RT SuperMix for qPCR (Vazyme Biotech Co..Ltd) was used to reverse transcribed RNA samples. RT-qPCR reactions were conducted with PerfectStart Green qPCR SuperMix (TransGen Biotech Co., Ltd). CFX Connect RealTime PCR Dection System (Bio-Rad Laboratories (Shanghai) Co., Ltd.) was used to detect the fluorescence and *scr* gene was selected to normalize sample data (zhou23). Each experiment was performed triplicate.

### Environmental L-alanine content assay

Measurement of environmental L-Alanine level was performed as the instruction of Amplite Fluorimetric L-Alanine Assay Kit (AAT Bioquest, Inc.). DSM media of sporulating cells at sporulation t_19_ was collected from the supernatant after centrifugation. The fluorescence intensity of L-Alanine in DSM media was monitored by TECAN Spark 10M microplate reader (TECAN, Switzerland) at Ex/Em = 540/590 nm. The standard solution of L-alanine was serially diluted and detected together to obtain a standard curve, and the L-Alanine content in the environment was calculated based on the standard curve.

### ATP content assay

Measurement of ATP content in mother cell was performed using the BacTiter-Glo Microbial Cell Viability Assay (Promega). As guided by the instructions, sporulating cells in DSM at sporulation t_1_ were collected and detected luminescence using a TECAN Spark 10M microplate reader (TECAN, Switzerland). The standard solution of ATP was serially diluted and detected simultaneously to obtain a standard curve, and the ATP level was calculated based on the standard curve.

### Data processing

Unless stated otherwise, each experiment was carried out at least triplicate. GraphPad Prism 8 software was used for all statistical analysis, data processing, and graph drawing. One-way ANOVA was performed to analyze the variance and p < 0.05 was regarded as significance for all data statistics.

### Data availability

The data that support the findings of this study are available from the authors on reasonable request.

## Acknowledgements

We are grateful to Dr. Bing Zhou (The Hebrew University of Jerusalem) for valuable discussions and comments. This work was supported by National Natural Science Foundation of China (NSFC) (grant No. 32372470), NSFC (grant No. 32001658), Agricultural Research Outstanding Talents of China (grant No. 13210317) awarded to Lei Rao, and 2115 Talent Development Program of China Agricultural University.

## Author contributions

F. L. and L. R. conceived the idea of the experiment. F. L. contributed to the acquisition and analysis of data as well as writing the draft. T. Z. contributed to the construction of mutant strains. Y. D., L. R., and X. L. contributed to revising the paper.

## Competing interests

The authors declare no competing interests.

## Notes

### Competing Interest Statement

The authors have declared no competing interest.

### Summary of Updates

We revised and updated the manuscript, mainly the discussion section. We supplemented the data in Figure 7 and 8, and added a new Figure 9.

