## Supplemental materials for "Alanine and glutamate catabolism ensure proper sporulation by preventing premature germination and providing energy respectively"

**Figure S1.** Representative phase-contrast images of sporulating cells at the late sporulation stage $t_{19}$. *B. subtilis* PY79 (wt), YZ24 (Δ*ald* Δ*rocG* Δ*gerAA, amyE::*$P_{IPTG}$*-rocG*), YZ25 (Δ*ald* Δ*rocG* Δ*gerAA, amyE::*$P_{IPTG}$*-ald*), and YZ26 (Δ*ald* Δ*rocG* Δ*gerAA, amyE::*$P_{IPTG}$*-rocG-ald*) strains were induced to sporulate in DSM at 37°C for 22 hrs and followed by microscopy. Different levels of IPTG were added to the YZ24, YZ25, and YZ26 cultures at the sporulation $t_0$ to induce corresponding gene expression. Scale bar, 2 μm.

**Figure S2.** Sporulation and germination phenotypes of mutant strains. (A) Quantification of YZ25 (Δ*ald* Δ*rocG* Δ*gerAA, amyE::*$P_{IPTG}$*-ald*) strains described in Figure S1. Data are presented as percentages of the number of the phase-bright spores and all sporulating cells in the same image ($n \geq 800$ for each strain); (B) AGFK-induced germination of spores collected in Figure S1. Spores of wt, as well as YZ24, YZ25, and YZ26 strains with (i) 10 μM and (ii) 20 μM IPTG induction, were purified and incubated with AGFK (10 mM) to trigger germination. DPA release was measured by detecting the relative fluorescence units (RFU) of $Tb^{3+}$-DPA. Shown is a representative experiment out of three independent biological repeats.

**Figure S3.** Cytological sporulation assay for Δ*rocG* mutants. Representative phase contrast and the indicated fluorescent images of wt and YZ19 (Δ*rocG*) cells harboring two transcriptional fusions, (A) $\sigma^F$ and (B) $\sigma^E$, at sporulation $t_2$, and $t_{2.5}$. Scale bar, 2 μm.

**Figure S1**

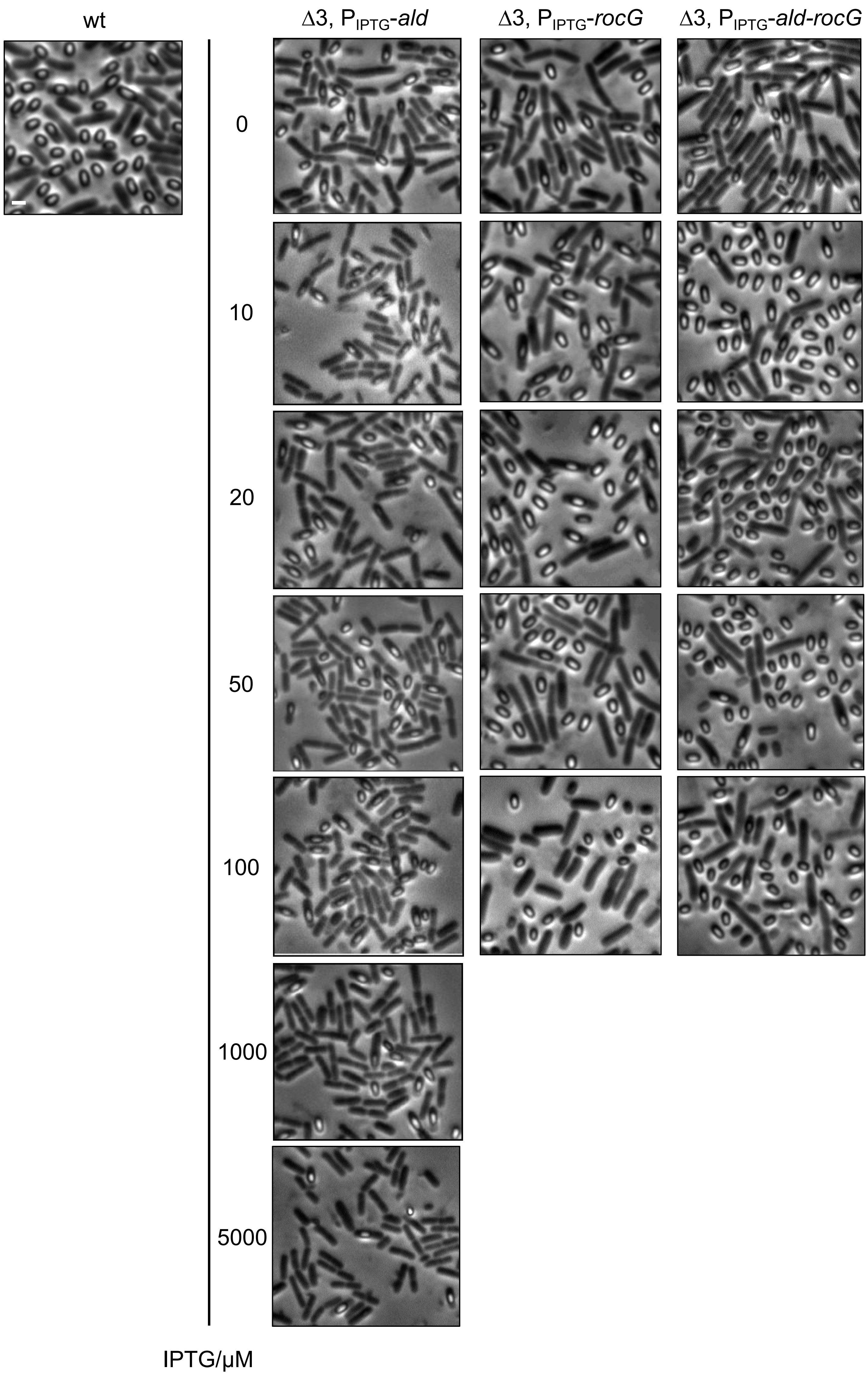
wt
Δ3, P$_{IPTG}$-*ald*
Δ3, P$_{IPTG}$-*rocG*
Δ3, P$_{IPTG}$-*ald-rocG*
0
10
20
50
100
1000
5000
IPTG/µM

### Figure S2

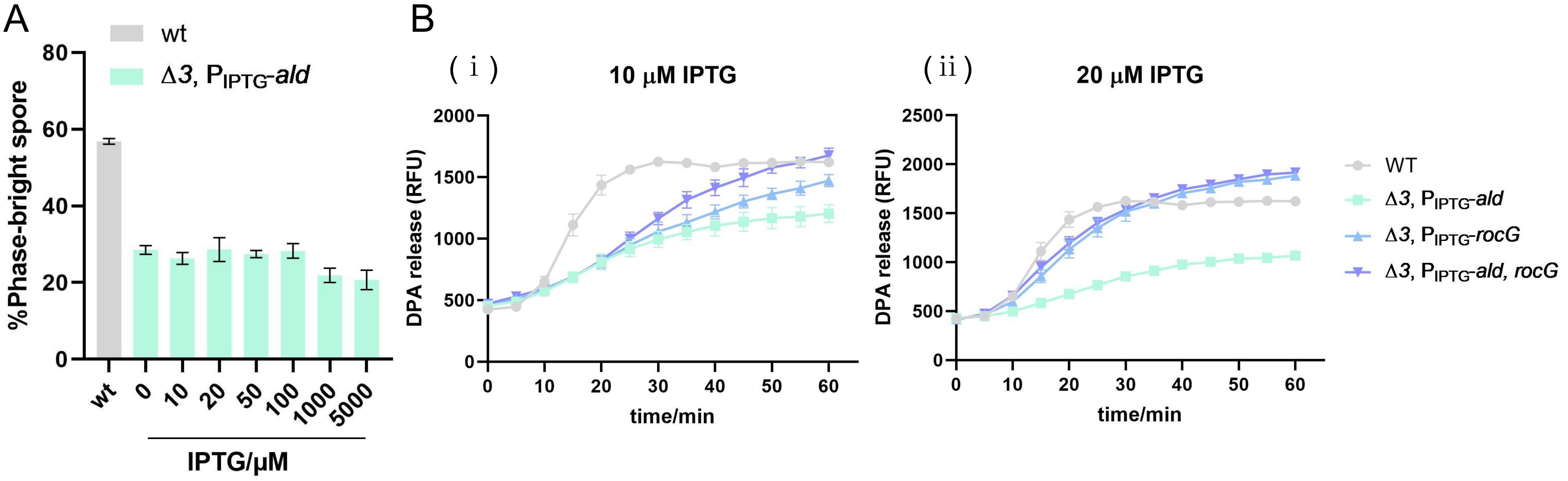
A
wt
Δ3, P_IPTG-ald
%Phase-bright spore
0
20
40
60
80
wt
0
10
20
50
100
1000
5000
IPTG/μM
B
( i )
10 μM IPTG
( ii )
20 μM IPTG
DPA release (RFU)
time/min
0
10
20
30
40
50
60
500
1000
1500
2000
2500
WT
Δ3, P_IPTG-ald
Δ3, P_IPTG-rocG
Δ3, P_IPTG-ald, rocG

**Figure S3**

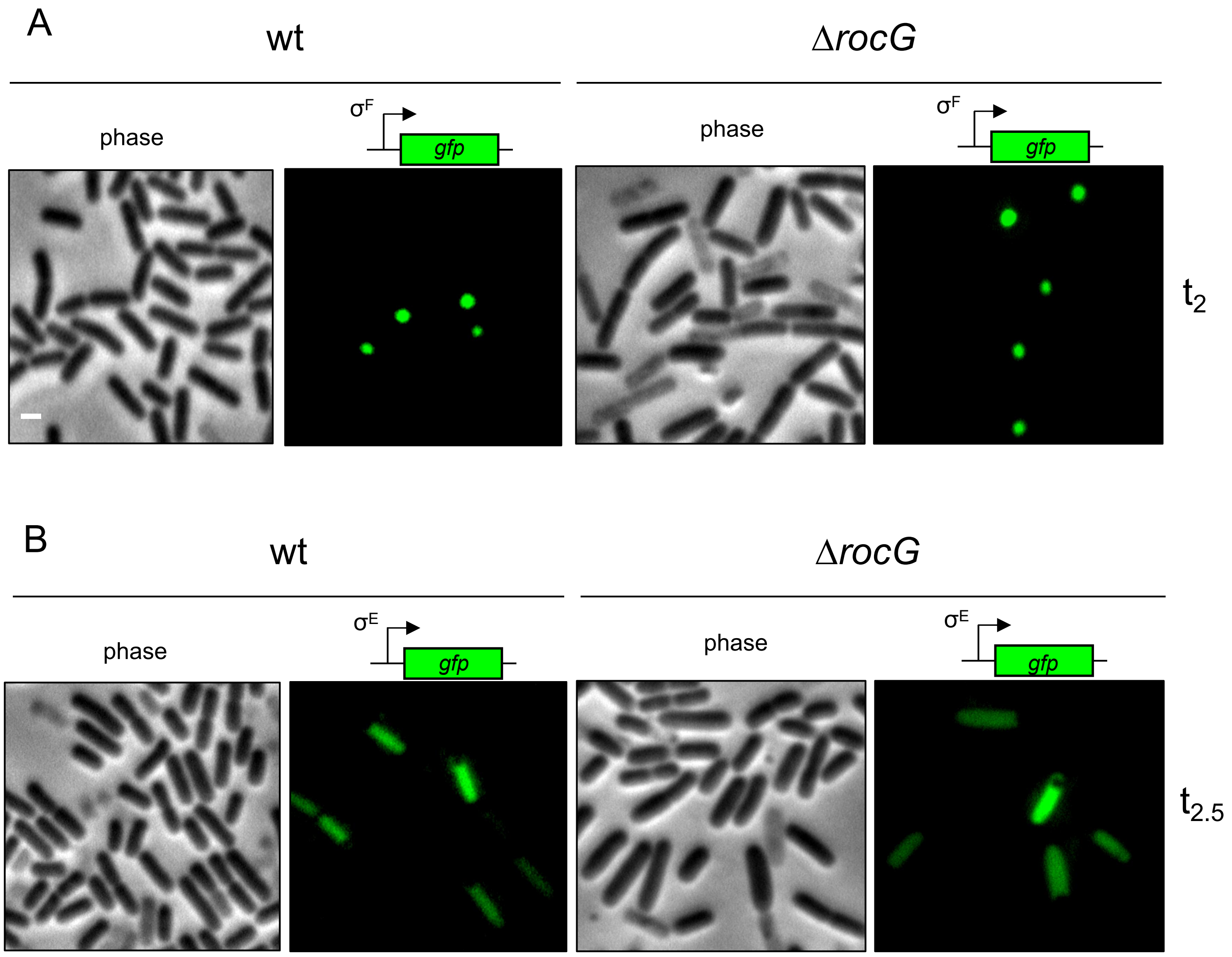
A
wt
ΔrocG
phase
σF
gfp
phase
σF
gfp
t2
B
wt
ΔrocG
phase
σE
gfp
phase
σE
gfp
t2.5

#### Table S1. Strains used in this study.

| Strain | Genotype | Source |
|---|---|---|
| PY79 | Wild type | Lab stock |
| LR32 | *rocR::mls, ahrC::kan* | Lab stock |
| YZ09 | *rocR::mls, ahrC::kan, adeR::cm* | This work |
| YZ10 | *gerAA::cm* | This work |
| YZ11 | *ald::mls* | This work |
| YZ12 | *ald::mls, rocG::kan* | This work |
| YZ13 | *ald::mls, rocG::kan, amyE::ald-rocG-cm* | This work |
| YZ18 | *adeR::cm* | This work |
| YZ19 | *rocG::kan* | This work |
| YZ21 | *ald::mls, gerAA::cm* | This work |
| YZ22 | *ald::mls, rocG::kan, gerAA::cm* | This work |
| YZ23 | *rocR::mls, ahrC::kan, gerAA::tet* | This work |
| YZ24 | *ald::mls, rocG::kan, gerAA::cm, amyE::*$P_{IPTG}$*-rocG-spc* | This work |
| YZ25 | *ald::mls, rocG::kan, gerAA::cm, amyE::*$P_{IPTG}$*-ald-spc* | This work |
| YZ26 | *ald::mls, rocG::kan, gerAA::cm, amyE::*$P_{IPTG}$*-rocG-ald-spc* | This work |
| YZ27 | *gerAA::tet* | This work |
| YZ31 | *ald::mls, amyE::*$P_{IPTG}$*-ald-spc* | This work |
| YZ32 | *rocG::kan, amyE::*$P_{IPTG}$*-rocG-spc* | This work |
| YZ81 | *adeR::cm, gerAA::tet* | This work |
| YZ90 | *rocR::mls, ahrC::kan, adeR::cm, gerAA::tet* | This work |
| BB01 | *amyE::*$P_{spoIIQ}$*-gfp-cm* | This work |
| BB02 | *amyE::*$P_{spoIID}$*-gfp-cm* | This work |
| BB03 | *amyE::*$P_{sspB}$*-gfp-cm* | This work |
| BB04 | *amyE::*$P_{gerE}$*-gfp-cm* | This work |
| BB09 | *rocG::kan, amyE::*$P_{spoIIQ}$*-gfp-cm* | This work |
| BB10 | *rocG::kan, amyE::* $P_{spoIID}$*-gfp-cm* | This work |
| BB11 | *rocG::kan, amyE::* $P_{sspB}$*-gfp-cm* | This work |
| BB12 | *rocG::kan, amyE::* $P_{gerE}$*-gfp-cm* | This work |

#### Detailed description of strain construction

YZ10 (*gerAA::cm)*: The ORF of *gerAA* was replaced by *cm* gene using long-flanking-homology PCR with primers oYZ101-oYZ104.

YZ11 (*ald::mls*): The ORF of *ald* was replaced by *mls* gene using long-flanking-homology PCR with primers oYZ111-oYZ114.

YZ19 (*rocG::kan*): PY79 was transformed with genomic DNA of strain BKK37790.

YZ12 (*ald::mls, rocG::kan*): YZ11 was transformed with genomic DNA of YZ19.

YZ13 (*ald::mls, rocG::kan, amyE::ald-rocG-cm*): YZ12 was transformed with pYZ13.

YZ21 (*ald::mls, gerAA::cm*): YZ11 was transformed with genomic DNA of YZ10.

YZ22 (*ald::mls, rocG::kan, gerAA::cm*): YZ12 was transformed with genomic DNA of YZ10.

YZ27 (*gerAA::tet*): The ORF of *gerAA* was replaced by *tet* gene using long-flanking-homology PCR with primers oYZ101-oYZ104.

YZ23 (*rocR::mls, ahrC::kan, gerAA::tet*): LR32 was transformed with genomic DNA of YZ27.

YZ24 (*ald::mls, rocG::kan, gerAA::cm, amyE::*$P_{IPTG}$*-rocG-spc*): YZ22 was transformed with pYZ14.

YZ25 (*ald::mls, rocG::kan, gerAA::cm, amyE::*$P_{IPTG}$*-ald-spc*): YZ22 was transformed with pYZ15.

YZ26 (*ald::mls, rocG::kan, gerAA::cm, amyE::*$P_{IPTG}$*-rocG-ald-spc*): YZ22 was transformed with pYZ16.

YZ31 (*ald::mls, amyE::*$P_{IPTG}$*-ald-spc*): YZ11 was transformed with pYZ15.

YZ32 (*rocG::kan, amyE::*$P_{IPTG}$*-rocG-spc*): YZ19 was transformed with pYZ14.

YZ18 (*adeR::cm*): The ORF of *adeR* was replaced by *cm* gene using long-flanking-homology PCR with primers oYZ181-oYZ184.

YZ09 (*rocR::mls, ahrC::kan, adeR::cm*): LR32 was transformed with genomic DNA of YZ18.

YZ81 (*adeR::cm, gerAA::tet*): YZ18 was transformed with genomic DNA of YZ27.

YZ90 (*rocR::mls, ahrC::kan, adeR::cm, gerAA::tet*): YZ09 was transformed with genomic DNA of YZ27.

BB01 (*amyE::*$P_{spoIIQ}$*-gfp-cm*): PY79 was transformed with pBB01.

BB02 (*amyE::*$P_{spoIID}$*-gfp-cm*): PY79 was transformed with pBB02.

BB03 (*amyE::* $P_{sspB}$*-gfp-cm*): PY79 was transformed with pBB03.

BB04 (*amyE::* $P_{gerE}$*-gfp-cm*): PY79 was transformed with pBB04.

BB09 (*rocG::kan, amyE::* $P_{spoIIQ}$*-gfp-cm*): YZ19 was transformed with pBB01.

BB10 (*rocG::kan, amyE::*$P_{spoIID}$*-gfp-cm*): YZ19 was transformed with pBB02.

BB11 (*rocG::kan, amyE::* $P_{sspB}$*-gfp-cm*): YZ19 was transformed with pBB03.

BB12 (*rocG::kan, amyE::* $P_{gerE}$*-gfp-cm*): YZ19 was transformed with pBB04.

#### Table S2. Plasmids used in this study.

| Plasmid | Description | Source |
|---|---|---|
| pDG364 | *amyE::cm* | Guerout-Fleury et al., 1996 (1) |
| pDR111 | *amyE::*$P_{IPTG}$*-spc* | a gift from David Rudner (Harvard Medical School) |
| pYZ13 | *amyE::ald-rocG-cm* | This work |
| pYZ14 | *amyE::*$P_{IPTG}$*-rocG-spc* | This work |
| pYZ15 | *amyE::*$P_{IPTG}$*-ald-spc* | This work |
| pYZ16 | *amyE::*$P_{IPTG}$*-rocG-ald-spc* | This work |
| pBB01 | *amyE::*$P_{spoIIQ}$*-gfp-cm* | This work |
| pBB02 | *amyE::*$P_{spoIID}$*-gfp-cm* | This work |
| pBB03 | *amyE::*$P_{sspB}$*-gfp-cm* | This work |
| pBB04 | *amyE::*$P_{gerE}$*-gfp-cm* | This work |

#### Detailed description of plasmid construction

pYZ13 (*amyE::ald-rocG-cm*): containing the *ald* gene (promoter and ORF) and *rocG* gene (promoter and ORF) with flanking *amyE* sequences and a *cm* gene, was constructed by amplifying the *ald* gene by PCR using primers oYZ131-oYZ132, the *rocG* gene by PCR using primers oYZ133-oYZ134. The PCR-amplified DNA was cloned into the BamHI site of pDG364 (*amyE::cm*) using Gibson Assembly Kit (NEB, USA).

pYZ14 (*amyE::*$P_{IPTG}$*-rocG-spc*): containing the *rocG* gene and IPTG inducible promoter with flanking *amyE* sequences and a *spc* gene, was constructed by amplifying the *rocG* gene by PCR using primers oYZ141 and oYZ142. The PCR-amplified DNA was cloned into the HindIII site of pDR111 (*amyE::*$P_{IPTG}$*-spc*) using Gibson Assembly Kit (NEB, USA).

pYZ15 (*amyE::*$P_{IPTG}$*-ald-spc*): containing the *ald* gene and IPTG inducible promoter with flanking *amyE* sequences and a *spc* gene, was constructed by amplifying the *ald* gene by PCR using primers oYZ151 and oYZ152. The PCR-amplified DNA was cloned into the HindIII site of pDR111 (*amyE::*$P_{IPTG}$*-spc*) using Gibson Assembly Kit (NEB, USA).

pYZ16 (*amyE::*$P_{IPTG}$*-rocG-ald-spc*): containing the *ald* gene, the *rocG* gene and IPTG inducible promoter with flanking *amyE* sequences and a spc gene, was constructed by amplifying the *rocG* gene by PCR using primers oYZ141 and oYZ161, the *ald* gene by PCR using primers oYZ162 and oYZ152. The PCR-amplified DNA was cloned into the HindIII site of pDR111 (*amyE::*$P_{IPTG}$*-spc*) using Gibson Assembly Kit (NEB, USA).

pBB01 (*amyE::*$P_{spoIIQ}$*-gfp-cm*): containing the *gfp* gene and *spoIIQ* promoter with flanking *amyE* sequences and a cm gene, was constructed by amplifying the *gfp* gene by PCR using primers oBB100 and oBB101, oBB011 and oBB012. The PCR-amplified DNA was cloned into the BamHI site of pDG364 (*amyE::cm*) using Gibson Assembly Kit (NEB, USA).

pBB02 (*amyE::P$_{spoIID}$-gfp-cm*): containing the *gfp* gene and *spoIID* promoter with flanking *amyE* sequences and a *cm* gene, was constructed by amplifying the *gfp* gene by PCR using primers oBB100 and oBB101, oBB021 and oBB022. The PCR-amplified DNA was cloned into the BamHI site of pDG364 (*amyE::cm*) using Gibson Assembly Kit (NEB, USA).

pBB03 (*amyE::P$_{sspB}$-gfp-cm*): containing the *gfp* gene and *sspB* promoter with flanking *amyE* sequences and a cm gene, was constructed by amplifying the *gfp* gene by PCR using primers oBB100 and oBB101, oBB031 and oBB032. The PCR-amplified DNA was cloned into the BamHI site of pDG364 (*amyE::cm*) using Gibson Assembly Kit (NEB, USA).

pBB04 (*amyE::P$_{gerE}$-gfp-cm*): containing the *gfp* gene and *gerE* promoter with flanking *amyE* sequences and a *cm* gene, was constructed by amplifying the *gfp* gene by PCR using primers oBB100 and oBB101, oBB041 and oBB042. The PCR-amplified DNA was cloned into the BamHI site of pDG364 (*amyE::cm*) using Gibson Assembly Kit (NEB, USA).

**Table S3. Primers used in this study.**

| Primer | Sequence |
|---|---|
| oYZ101 | TAGCATCCTGAAGATGCGTG |
| oYZ102 | CTGAGCGAGGGAGCAGAATGAGGTCACCTCTTATCCAAAACCTTAG |
| oYZ103 | GTTGACCAGTGCTCCCTGTACCTTTCAAGGTATTTCTATTGTTGCA |
| oYZ104 | GATAATCACCATGCTTGTTTTAGTCGATA |
| oYZ111 | TTCATATGGGCGCTTGAGAT |
| oYZ112 | CTGAGCGAGGGAGCAGAAATCTGTCTCCTCCTGTATATGTG |
| oYZ113 | GTTGACCAGTGCTCCCTGTTCACAATAAGCTTGCAGAAAGATTTC |
| oYZ114 | AATGAAGAAGGCCAATATTCCCTAT |
| oYZ181 | AATATCAATTACTTTTTTCACCGGGTGG |
| oYZ182 | CTGAGCGAGGGAGCAGAACTCATCCCCTTCCTTTTATTATGATTGT |
| oYZ183 | GTTGACCAGTGCTCCCTGAATTTCACAAATCCATGTTTTTTTATTTTCT<br>TAATC |
| oYZ184 | TTTCATATGTTGGCTGATCATGTGTT |
| oYZ131 | CCAACTGGTAATGGTAGCGACCGGCGCTCAG<br>GGATCAGCCAGATCGCTGAAATT |
| oYZ132 | GGGGATCAATCACCAGCTG TTAAGCACCCGCCACAGAT |
| oYZ133 | CAGCTGGTGATTGATCCCC |
| oYZ134 | GTCAAACATGAGAATTCGATAAGCTTCTAG<br>ATATCTTCGCATATGAATAAAAATGCAAGAATCATGCC |
| oYZ141 | AATTGTGAGCGGATAACAATTAAGCTTAAAGGTGGTGAACTACT<br>ATCCAATGAA GAAAGGAATTGGC |
| oYZ142 | ATGCGGCTAGCTGTCGACTATCAAAAATGCAAGAATCATGCCATTTTCT |
| oYZ151 | AATTGTGAGCGGATAACAATTAAGCTT<br>AAAGGTGGTGAACTACTATGATCATAGGGGTTCCTAAAGAGA |
| oYZ152 | ATGCGGCTAGCTGTCGACTATCATTAAGCACCCGCCACAG |
| oYZ161 | AAAATGCAAGAATCATGCCATTTTCT |
| oYZ162 | AGAAAATGGCATGATTCTTGCATTTT<br>ATGATCATAGGGGTTCCTAAAGAGA |
| oBB100 | ATGAGTAAAGGAGAAGAACTTTTCACTG |
| oBB101 | GTCAAACATGAGAATTCGATAAGCTTCTAGTCA<br>TTTGTATAGTTCATCCATGCCATGTG |
| oBB011 | CCAACTGGTAATGGTAGCGACCGGCGCTCAG<br>AACAGCCTCCAAGTCTCTATG |
| oBB012 | CAGTGAAAAGTTCTTCTCCTTTACTCAT<br>TGTTTCATCACCTCAGCAACATT |
| oBB021 | CCAACTGGTAATGGTAGCGACCGGCGCTCAG<br>GCGCAAAATAGCAAAA AAGAATACGT |
| oBB022 | CAGTGAAAAGTTCTTCTCCTTTACTCAT<br>ATTCAGCTGCCTCCTGCT |
| oBB031 | CCAACTGGTAATGGTAGCGACCGGCGCTCAG |

| | |
|---|---|
| | CTTAGCCTAAACGGCTAAGC |
| oBB032 | CAGTGAAAAGTTCTTCTCCTTTACTCAT TTGCACCAATCTGTGAACC |
| oBB041 | CCAACTGGTAATGGTAGCGACCGGCGCTCAG AAACGTCACCTCCTGCGC |
| oBB042 | CAGTGAAAAGTTCTTCTCCTTTACTCAT GTATTGTAACCCTCCTTGCTAAGGTG |
